# A faster and more accurate algorithm for calculating population genetics statistics requiring sums of Stirling numbers of the first kind

**DOI:** 10.1101/2020.03.12.989392

**Authors:** Swaine L. Chen, Nico M. Temme

## Abstract

Stirling numbers of the first kind are used in the derivation of several population genetics statistics, which in turn are useful for testing evolutionary hypotheses directly from DNA sequences. Here, we explore the cumulative distribution function of these Stirling numbers, which enables a single direct estimate of the sum, using representations in terms of the incomplete beta function. This estimator enables an improved method for calculating an asymptotic estimate for one useful statistic, Fu’s *F_s_*. By reducing the calculation from a sum of terms involving Stirling numbers to a single estimate, we simultaneously improve accuracy and dramatically increase speed.

## 1 Introduction

The dominant paradigm in population genetics is based on a comparison of observed data with parameters derived from a theoretical model [1, 2]. Specifically for DNA sequences, many techniques have been developed to test for extreme relationships between average sequence diversity (number of DNA differences between individuals) and the number alleles (distinct DNA sequences in the population). In particular, such methods are widely used to predict selective pressures, where certain mutations confer increased or decreased survival to the next generation [2]. Such selective pressures are relevant for understanding and modeling practical problems such as influenza evolution over time [3] and during vaccine production [4]; adaptations in human populations, which may impact disease risk [5, 6]; and the emergence of new infectious diseases and outbreaks [7].

Many population genetics tests are therefore formulated as unidimensional test statistics, where the pattern of DNA mutations in a sample of individuals is reduced to a single number [2, 1, 8]. Such statistics are heavily informed by combinatorial sampling and probability distribution theories, many of which are built upon the foundational Ewens’s sampling formula [9]. Ewens’s sampling formula describes the expected distribution of the number of alleles in a sample of individuals, given the nucleotide diversity. Calculation of subsets of this distribution are useful for testing deviations of observed data from a null model; such subsets often require the calculation of Stirling numbers of the first kind (hereafter referred to simply as Stirling numbers). In particular, two population genetics statistics, the Fu’s *F_s_* and Strobeck’s *S* statistics, utilize this approach [8, 10]. The former has recently been shown to be potentially useful for detecting genetic loci under selection during population expansions (such as an infectious outbreak) both in theory and in practice [7]. However, Stirling numbers rapidly grow large and overwhelm the standard floating point range of modern computers.

In previous work, an asymptotic estimate for individual Stirling numbers was used to solve the problem of computing Fu’s *F_s_* for large datasets that are now becoming common due to rapid progress in DNA sequencing technology [11]. This new algorithm solved problems of numerical overflow and underflow, maintained good accuracy, and substantially increased speed compared with other existing software packages. However, the estimation of individual Stirling numbers led to the use of an estimator at least *n* – *m* +1 and at most 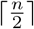 times. Here, we explore the potential for further increasing both accuracy and speed in calculating Fu’s *F_s_* by using a single estimator.

The new estimator for Fu’s *F_s_* has been implemented in R and is available at https://github.com/swainechen/hfufs.

## 2 Background Theory

### 2.1 General definitions

We take a population of *n* individuals, each of which carries a particular DNA sequence *D_i_* (referred to as the allele of individual *i*). We define a metric, dist(*D_i_, D_j_*) to be the number of positions at which sequence *D_i_* differs from *D_j_*. Then, we denote the average pairwise nucleotide difference as *θ_π_* (hereafter referred to simply as *θ*), defined as:

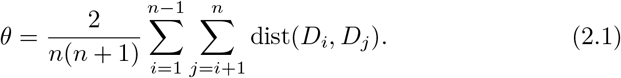

We also define a set of unique alleles *Du_i_* ∈ {*D_i_*} which have the property of 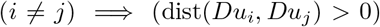. The ordinality of {*Du_i_*} is denoted *m*, i.e. the number of distinct alleles in the data set.

Building upon on Ewens’s sampling formula [8, 9], it has been shown that the probability that, for given *n* and *θ*, at least *m* alleles would be found, is

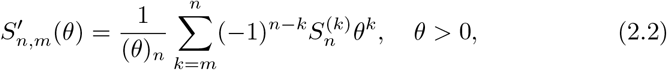

where (*θ*)_*n*_ is the Pochhammer symbol, defined by

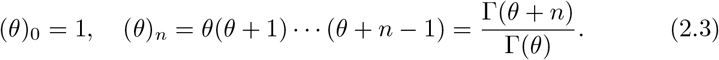

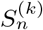 is a Stirling number and is defined by:

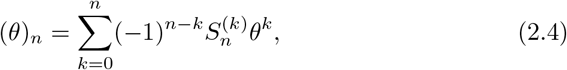

Fu’s *F_s_* is then defined as:

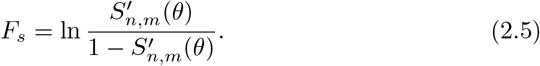

Fu’s *F_s_* thus measures the probability of finding a more extreme (equal or higher) number of alleles than actually observed. It requires computing a sum of terms containing Stirling numbers, which rapidly become large and therefore impractical to calculate explicitly even with modern computers [11].

Because of the relation in (2.4), the statistics quantity 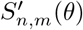 satisfies 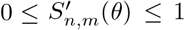. Also, this relation and (2.3) show that 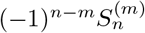 are nonnegative. We have the special values

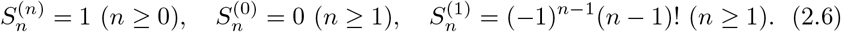

There is a recurrence relation

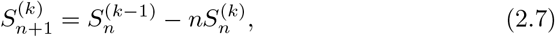

which easily follows from (2.4). For a concise overview of properties, with a summary of the uniform approximations, see [12, §11.3].

We introduce a complementary relation

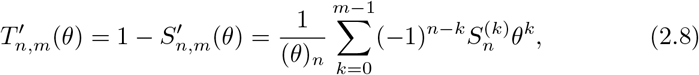

leading to an alternate calculation for Fu’s *F_s_* of

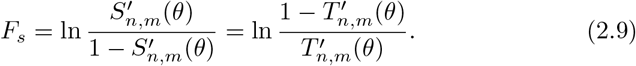

The recent algorithm considered in [11] is based on asymptotic estimates of 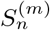 derived in [13], which are valid for large values of *n*, with unrestricted values of *m* ∈ (0, *n*). It avoids the use of the recursion relation given in (2.7).

In the present paper we derive an integral representation of 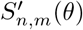 and of the complementary function 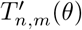, for which we can use the same asymptotic approach as for the Stirling numbers without calculating the Stirling numbers themselves. From the integral representation we also obtain a representation in which the incomplete beta function occurs as the main approximant. In this way we have a convenient representation, which is available as well for many classical cumulative distribution functions. We show numerical tests based on a first- order asymptotic approximation, which includes the incomplete beta function. In a future paper we give more details on the complete asymptotic expansion of 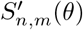, and, in addition, we will consider an inversion problem for large *n* and *m*: to find *θ* either from the equation 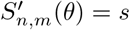, when *s* ∈ (0,1) is given, or from the equation *F_s_* = *f*, when *f* ∈ ℝ is given.

### 2.2 Remarks on computing 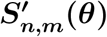

When computing the quantity *F_s_* defined in (2.5), numerical instability may happen when 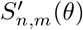 is close to 1. In that case, the computation of 1 – *S*’ suffers from cancellation of digits. For example, take *n* = 100, *θ* = 39.37, *m* = 31. Then 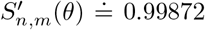, and *F_s_* becomes about 6.6561 when using the first relation in (2.9). However, when we calculate 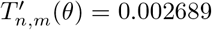 and use the second relation, then we obtain the more reliable result *F_s_* =. 5.9160.

We conclude that, when 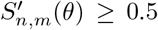, it is better to switch and obtain 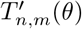 from the sum in (2.8), and by using the second relation of *F_s_* in (2.9). A simple criterion to decide about this can be based on using the saddle point *z*_0_ (see Remark 3.1 below).

A second point is the overflow in numerical computations when *n* is large, because of the large values of 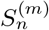 when *m* is small with respect to *n*. For example, when *n* = 10, *m* = 5 we have

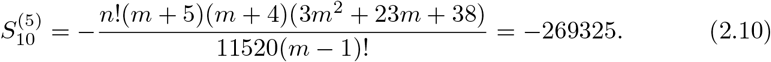

Therefore, it is convenient to scale the Stirling number in the form 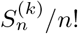. In addition, the Pochhammer term (*θ*)*n* in front of the sum in (2.2) will also be large with n; we have (1)_*n*_ = *n*!.

We can write the sum in (2.2) in the form

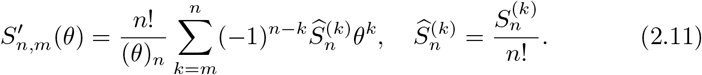

Leading to a corresponding modification in the recurrence relation in (2.7) for the scaled Stirling numbers:

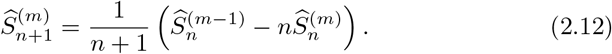

To control overflow, we can consider the ratio

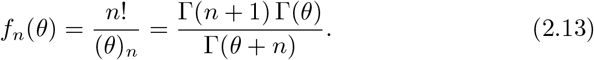

This function satisfies *f_n_*(*θ*) ≤ 1 if *θ* ≥ 1. For small values of *n* we can use recursion in the form

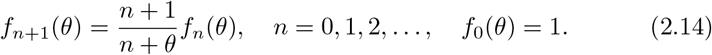

For large values of *n* and all *θ* > 0 we can use a representation based on asymptotic forms of the gamma function.

#### Remark 2.1.

It should be observed that using the recursion in (2.7) and (2.12) is a rather tedious process when *n* is large. For example, when we use it to obtain 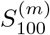 for all *m* ∈ (0, 100], we need all previous 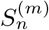 with *n* ≤ 99 for all *m* ∈ (0, *n*]. Table look-up for 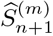 in floating point form may be a solution. When *n* is large enough, the algorithm mentioned in [11] evaluates each needed Stirling number by using the asymptotic approximation derived in [13].

## 3 Results and Discussion

### 3.1 An integral representation of 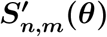

We use the integral representation of the Stirling numbers that follows from the definition given in (2.4). That is, by using Cauchy’s formula,

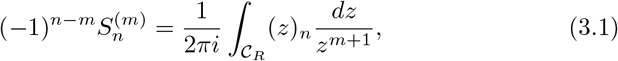

where 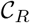 is a circle around the origin with radius *R*. We can take *R* as large as we like. As in [13, §3], it is convenient to proceed with

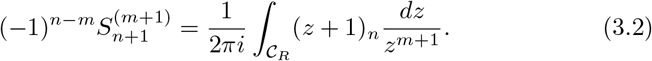

We derive an integral representation of

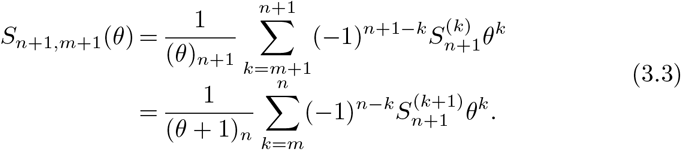

We use (3.2) and obtain

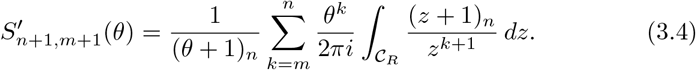

We can take *R* > *θ* to have |*θ/z*| < 1 on the circle 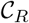, and we can perform the summation to ∞, because all terms with *k* > *n* do not give contributions. In this way we obtain the requested integral representation

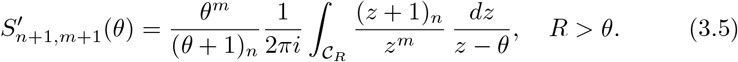

To obtain this result we need *R* > *θ*, but in the integral representation we can take *R* < *θ* when we pick up the residue at *z* = *θ*. The result is

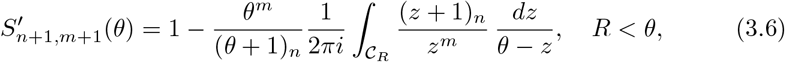

and we find for 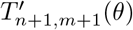 (see (2.8))

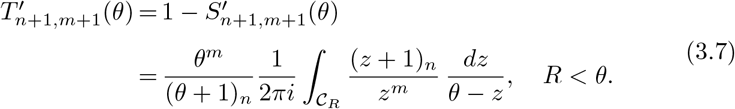

For the asymptotic analysis we write (3.5) in the form

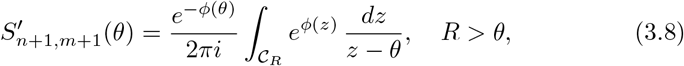

where

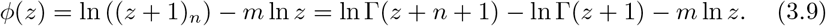

Then the saddle point of the integral in (3.8) follows from the equation

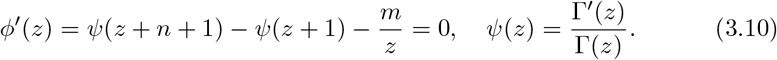

There is a positive saddle point *z*_0_ when 0 < *m* < *n*.

#### Remark 3.1.

When *θ* crosses the value 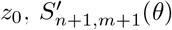 becomes (almost) 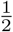. Especially when the parameters *m* and *n* are large, 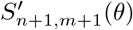 starts with very small values for small *θ*, its values is about 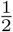 when *θ* = *z*_0_ and it becomes quickly 1 as *θ* increases. We call *z*_0_ the transition value for *θ*.

For fixed values of *θ* there is also a transition value for *m*, say, *m*_0_. When *n* is large, 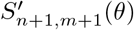 starts at values near 1 for small *m*, it becomes about 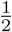 when *m* crosses the transition value *m*_0_, and it becomes quickly small as *m* → *n*.

### 3.2 An asymptotic representation of 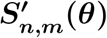

We use the transformation, as in [13, §3],

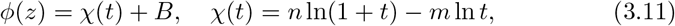

with condition sign(*z* – *z*_0_) = sign(*t* – *t*_0_), where *t*_0_ is the saddle point in the *t*-domain and also the zero of

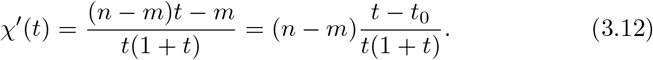

Also

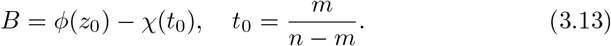

With this choice of *B*, the variables *z* and *t* correspond with each other at the respective saddle points.

Using the transformation we obtain

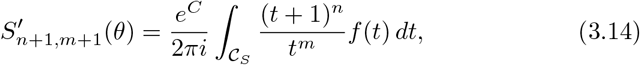

where

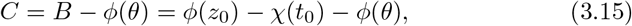

and

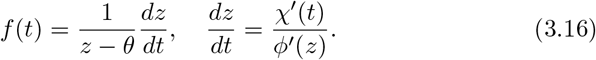

The contour 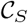 runs around the origin and includes a pole at *t* = *τ* that corresponds with the pole in the *z*-plane at *z* = *θ*.

### 3.3 A representation in terms of the incomplete beta function

The integrands of the integral representations of 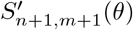 have a pole at *z* = *θ*. For the contour integrals this is not a complication, because by using the theory of analytic functions we can deform the contour to avoid the pole, and we can even cross the pole and pick up the residue as we did to obtain the representation in (3.6).

For the integral in the *t*-domain given in (3.14) the same can be done. The function *f*(*t*) has a pole at a point *t* = *τ*, say, that follows from the transformation given in (3.11). That means, *τ* is defined by the equation

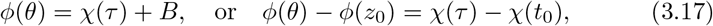

and we can show the existence of the pole of the function *f*(*t*) defined in (3.16) writing

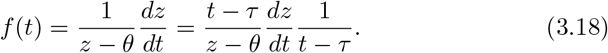

In asymptotic analysis the presence of such a pole is of great interest, especial when (in the *t*-domain) the saddle point (here *t*_0_) is close to a pole (here *τ*), or even when these points coalesce. See, for example, [14, Chapter 21]. Usually, the error function is introduced to handle the asymptotic analysis, in the present can we use an incomplete beta function.

We split off the pole from *f*(*t*) and write

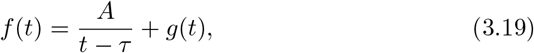

where we assume that *g*(*t*) is well defined at *t* = *τ*. To find *A* we use the analytical relation in (3.11) between *t* and *z*, in particular at *z* = *θ* (or *t* = *τ*). Applying l’Hôpital’s rule, we conclude that 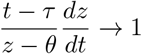 as *t* → *τ*, which gives *A* = 1. Hence, substituting this form of *f*(*t*) in (3.14), we find

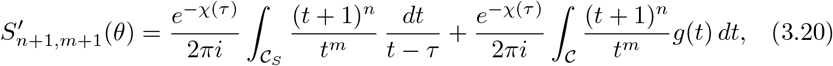

where we have used (see (3.15) and (3.17))

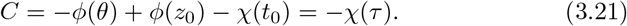

The radius of the circle 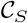 in the first integral is larger than *τ*, for the second integral we take a circle 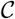 around the origin such that the singularities of *g*(*t*) are outside the circle.

The first integral can be evaluated in terms of the incomplete beta function defined by

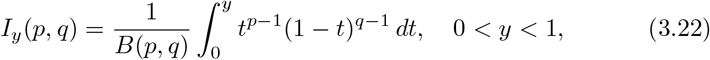

where *B*(*p, q*) is the complete beta function

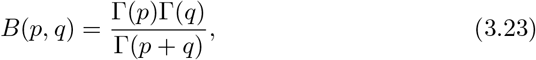

We will show in the Appendix that

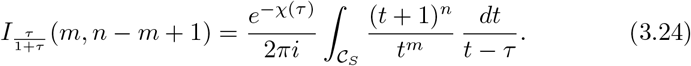

Hence,

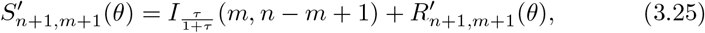

where

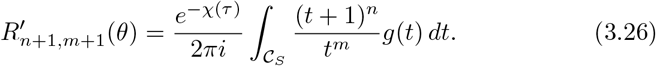

A first-order approximation of this function follows from replacing *g*(*t*) by its value at the saddle point *t*_0_. This gives

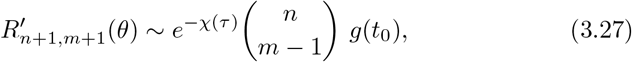

where

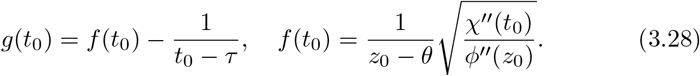

This expression of *f*(*t*_0_) follows from the definition of *f*(*t*) given in (3.16). In a future publication we will give details about the complete asymptotic expansion of the term 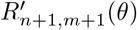.

For the complementary function (see (2.8)) we obtain

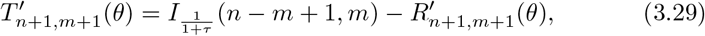

where we have used

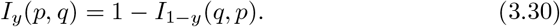

#### Remark 3.2.

The incomplete beta function in (3.25) has the representation (see [15, §8.17(i)])

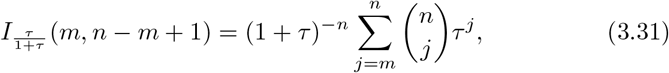

and from the complementary relation in (3.30) it follows that the function in (3.29) has the expansion

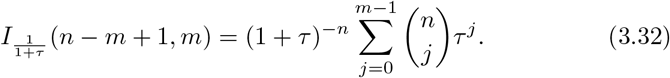

### 3.4 Numerical tests

We summarize the steps to obtain the first-order approximations (see (3.25) or (3.29) and (3.27))

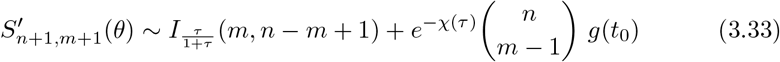

or

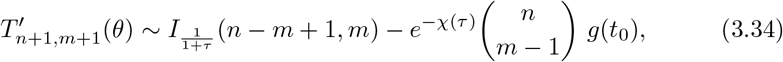

for given *θ, n* and *m*, and to compute Fu’s *F_s_* by using (2.9).

1. Compute the saddle point *z*_0_, the positive zero of *ϕ*′(*z*); see (3.10).
2. With *t*_0_ = *m*/(*n* – *m*), the positive zero of *χ*’(*t*) (see (3.12)), compute *τ*, the solution of the equation (see (3.17))

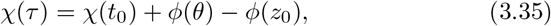

with *ϕ*(*z*) defined in (3.9) and *χ*(*t*) defined in (3.11). When *θ* = *z*_0_ there is one solution *τ* = *t*_0_. When *τ* = *t*_0_ there are two positive solutions, and we take the one that satisfies the condition sign(*θ* – *z*_0_) = sign(*τ* – *t*_0_).
3. When *θ* < *z*_0_, hence *τ* < *t*_0_, compute the approximation of 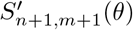 by using (3.33), and *F_s_* from the first relation in (2.9).
4. When *θ* > *z*_0_, hence, *τ* > *t*_0_, compute the approximation of 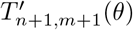 by using (3.34), and *F_s_* from the second relation in (2.9).

In Table 1 we give the relative errors in the computation of *F_s_* defined in (2.5). The values of *n, m*, and *θ* correspond with those in Table 1 of [11]. We have used the asymptotic result (3.27). Computations are done with Maple, with Digits = 16. The “exact” values are obtained by using Maple’s code for Stirling1 (n, m), which computes the Stirling numbers of the first kind.

**Table 1:** Relative errors in the computation of *F_s_* defined in (2.5). We have used the asymptotic result (3.27).

| $n/m$ | $\theta$ | $F_s$, asymptotic | $F_s$, exact | rel.error |
| --- | --- | --- | --- | --- |
| 25/20 | 9.39 | -6.83168 | -6.8294578 | $0.33 \times 10^{-3}$ |
| 50/31 | 9.61 | -10.13052 | -10.1290263 | $0.15 \times 10^{-3}$ |
| 100/40 | 9.37 | -10.23064 | -10.2298131 | $0.81 \times 10^{-4}$ |
| 250/67 | 8.96 | -26.41607 | -26.4155959 | $0.18 \times 10^{-4}$ |
| 500/95 | 9.04 | -46.76268 | -46.76238956 | $0.63 \times 10^{-5}$ |
| 1000/152 | 9.07 | -112.42500 | -112.4248080 | $0.17 \times 10^{-5}$ |
| 2001/213 | 9.03 | -192.21835 | -192.2182390 | $0.60 \times 10^{-6}$ |

We additionally performed a comparison with the recently published algorithm in [11]. We performed 10,000 calculations with each algorithm and compared the results with an exact calculator. As expected, since the previous algorithm required estimating a Stirling number for each term of the sum, while the current asymptotic estimate directly calculates the sum, both error and compute speed were improved. Relative error for the single term estimate in (3.25) was well controlled at < 0.001 for nearly 99% of the calculations; for 411 calculations where the previous hybrid estimator had an error > 0.001, the estimate in (3.25) was more accurate in all but one case (*n* = 157, *m* = 4, *θ* = 43.59732; 3.08e-3 relative accuracy using [11]; 3.32e-3 relative accuracy using (3.25)) (Figure 1). The fewer calculations led to a clear improvement in calculation speed (median 54.6x faster; Figure 2).

**Figure 1:**
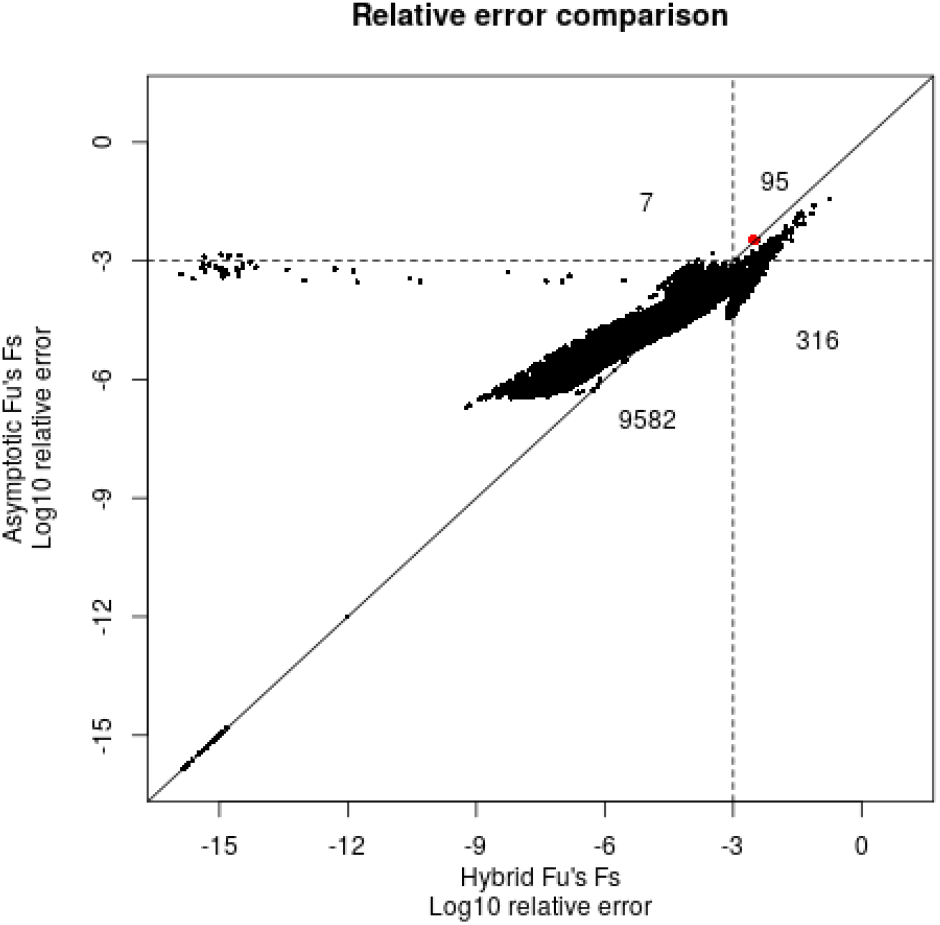
Comparison of relative error of the estimator from [11] and the single term asymptotic estimator in (3.25). Relative error for each is calculated against the arbitrary precision implementation described in [11]. In total, 10,000 calculations were performed with *n* randomly sampled from a uniform distribution between 50 and 500; *m* between 2 and *n*; and *θ* between 1 and 50. A solid diagonal line is drawn at *y* = *x*. Dotted lines are drawn at a relative error of 0.001. Numbers within each quadrant defined by the dotted lines indicate the number of points in each quadrant. The red dot indicates the one case where the relative error was > 0.001 and the error of (3.25) was greater than the estimator from [11].

**Figure 2:**
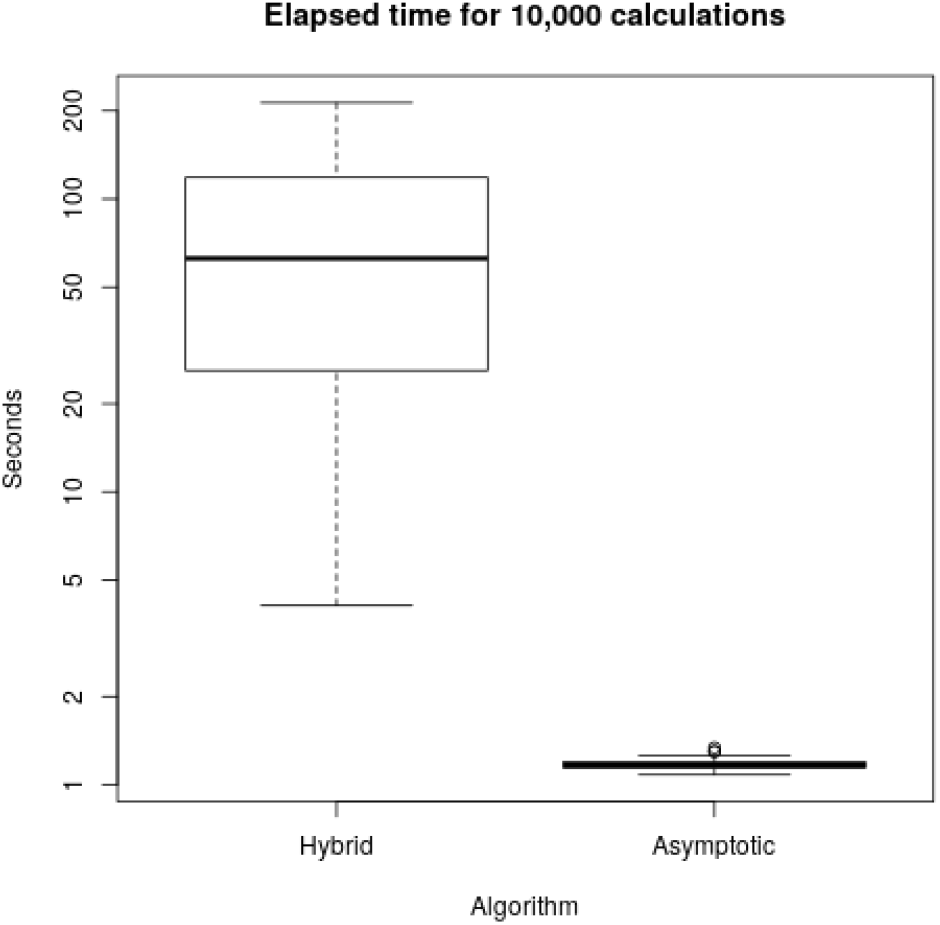
Comparison of run times between the hybrid algorithm from [11] and the single term asymptotic estimator in (3.25). 100 iterations were run, each with 10,000 calculations. The same set of parameters were used for each algorithm. The order of running the algorithms was alternated with each iteration. The dark horizontal line indicates the median, the box indicates the first and third quartiles, the whiskers are drawn at 1.5x the interquartile range, and outliers are represented by open circles. The median for the hybrid algorithm is 62.64 s; the median for the asymptotic algorithm is 1.17 s.

## 4 Conclusion

The rapid growth of sequencing data has been an enormous boon to population genetics and the study of evolution. Traditional population genetics statistics are still in common use today. The statistics Fu’s *F_s_* and Strobeck’s *S* have been difficult to calculate using previous methods; we now further improve both accuracy and speed for the calculation of Fu’s *F_s_* for large, modern data sets, using the main estimator in (3.25). Our plan for a paper about the ability to invert the calculation provides additional future directions in understanding the performance of these statistics, and the methods used herein may be useful for the development of new statistics that more effectively capture different types of selection.

## Acknowledgments

SLC acknowledges Shyam Prabhakar and members of the Chen lab for fruitful discussions. NMT acknowledges CWI, Amsterdam, for scientific support.

SLC was supported by the National Medical Research Council, Ministry of Health, Singapore (grant numbers NMRC/OFIRG/0009/2016 and NMRC/CIRG/1467/2017).

NMT was supported by the Ministerio de Ciencia e Innovaci’on, Spain, projects MTM2015-67142-P (MINECO/FEDER, UE) and

PGC2018-098279-B-I00 (MCIU/AEI/FEDER, UE). The authors affirm that all data necessary for confirming the conclusions of the article are present within the article, figures, and tables.

## 5 Appendix

We give a proof of the incomplete beta integral in (3.17). We use the integral representation of the hypergeometric function (see [16, §15.6])

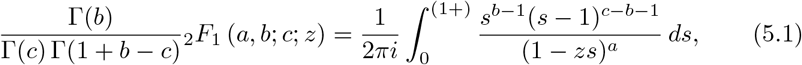

where the contour starts at the origin, encircles the point *s* = 1 in the anticlockwise direction, and returns to the origin. The main conditions are 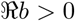 and that *s* = 1 /*z* is outside the contour.

We also use the relation between the incomplete beta function and the _2_*F*_1_-function (see [15, §8.17(ii)])

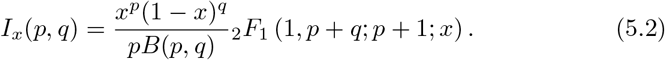

It follows that

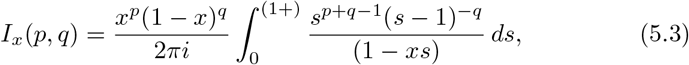

and after the substitution *s* = 1 + *t* and writing *x* = 1/(1 + *τ*), we obtain with *p*=*n*–*m*+1, *q*=*m*, and *χ*(*τ*) asin (3.11)

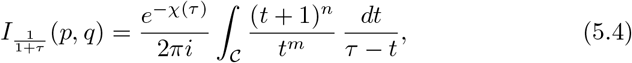

where the pole at *t* = *τ* is outside the contour. We modify the contour and pick up the residue. In this way we find the relation in (3.17).

